# Retinotopic organization of visual cortex in human infants

**DOI:** 10.1101/2020.12.01.407437

**Authors:** C. T. Ellis, T. S. Yates, L. J. Skalaban, V. R. Bejjanki, M. J. Arcaro, N. B. Turk-Browne

**Affiliations:** Department of Psychology, Yale University, New Haven, CT 06511, USA; Department of Psychology, Hamilton College, Clinton, NY 13323, USA; Department of Psychology, University of Pennsylvania, Philadelphia, PA 19104, USA

## Abstract

Vision develops rapidly during infancy, yet how visual cortex is organized during this period is unclear. One possibility is that the retinotopic organization of visual cortex emerges gradually as perceptual abilities improve. This may result in a hierarchical maturation of visual areas from striate to extrastriate cortex. Another possibility is that retinotopic organization is present from early infancy. This early maturation of area boundaries and tuning could scaffold further developmental changes. Here we test the functional maturity of infant visual cortex by performing retinotopic mapping with fMRI. Infants aged 5–23 months had retinotopic maps, with alternating preferences for vertical and horizontal meridians indicative of area boundaries from V1 to V4, and an orthogonal gradient of preferences from high to low spatial frequencies indicative of growing receptive field sizes. Although present in the youngest infants, these retinotopic maps showed subtle agerelated changes, suggesting that early maturation undergoes continued refinement.

## Introduction

Vision is the dominant sense in humans, but develops slowly throughout childhood and even into adolescence (Braddick and Atkinson, 2011; Kiorpes, 2016; Lewis and Maurer, 2005). How is infant visual cortex organized and how does this organization change over early development? One hypothesis is that visual cortex develops hierarchically (Bourne and Rosa, 2006), with low-level areas (such as V1) maturing first, followed by mid-level areas (such as V2–V4), and then high-level areas (such as LO and PHC) (Bourne and Rosa, 2006; Condé, Lund, and Lewis, 1996; Distler, Bachevalier, Kennedy, Mishkin, and Ungerleider, 1996; Gomez, Natu, Jeska, Barnett, and Grill-Spector, 2018; Kiorpes, 2016; Zhang, Zheng, Watanabe, Maruko, Bi, Smith, and Chino, 2005). By maturation, we consider the emergence of an organized retinotopic map of visual space that defines the tuning and boundaries of an area (i.e., arealization). According to hierarchical maturation, young infants may have mature arealization in only V1, whereas older infants may additionally show arealization in V2–V4. An alternative hypothesis is that the organization of visual cortex is established early in infancy. By this early arealization account, even young infants may have distinct retinotopic maps in areas V1–V4 with stereotyped tuning to features such as curvature and scale (Arcaro and Livingstone, 2017).

In support of hierarchical maturation, animal models show a sequence of cellular and macroscopic changes across the cortical hierarchy (Condé, Lund, and Lewis, 1996; Distler, Bachevalier, Kennedy, Mishkin, and Ungerleider, 1996; Zhang and Fang, 2012). At birth, mature cells are only found in low-level visual areas, whereas in the weeks following birth, they can be found through-out the visual hierarchy (Bourne and Rosa, 2006), including mid- and high-level areas. The behavioral capacities of humans similarly suggest sequential development of visual areas (Kovács, 2000; Siu and Murphy, 2018). Visual behaviors thought to rely primarily on V1, such as orientation discrimination and spatial frequency discrimination (Banks, Stephens, and Hartmann, 1985; Braddick, Wattam-Bell, and Atkinson, 1986), are present in rudimentary form near birth. More complex visual behaviors thought to depend on V2–V4 and interconnectivity between visual areas (Burkhalter, Bernardo, and Charles, 1993; Kiorpes and Bassin, 2003; Zhang, Zheng, Watanabe, Maruko, Bi, Smith, and Chino, 2005), such as contour integration (Baker, Tse, Gerhardstein, and Adler, 2008; Kovacs, Kozma, Feher, and Benedek, 1999), develop up to a year later. Indeed, the receptive field properties of high-level visual areas continue to develop during childhood, whereas those of low- and mid-level visual areas do not (Gomez, Natu, Jeska, Barnett, and Grill-Spector, 2018). In support of the early arealization hypothesis, arealization is present throughout the visual hierarchy in neonatal macaques (Arcaro and Livingstone, 2017). At 2 weeks old — homologous to 2 months old in humans (Boothe, Dobson, and Teller, 1985; Kiorpes, 2016) — functional connectivity in macaque visual cortex mirrors boundaries between retinotopic areas. Although these two hypotheses about arealization are juxtaposed here, they are not necessarily mutually exclusive: the boundaries of areas may be present early in infancy with other properties, such as receptive field size, maturing at different rates across areas.

The question of how visual cortex is organized in human infants remains unanswered because it has not been studied directly. Current theorizing is limited because it depends upon neural data from non-human animals and behavioral data from humans. For example, evidence for early arealization in macaques might not translate to humans. Even though the visual systems of humans and macaques are similar at maturity (Wandell, Dumoulin, and Brewer, 2007), human infancy is more protracted than macaque infancy (Boothe, Dobson, and Teller, 1985; Kiorpes, 2016). This allows for the possibility that retinotopic maps emerge in the human brain over postnatal development, consistent with hierarchical maturation. Likewise, the sequential emergence of visual behaviors in human infants could reflect hierarchical maturation of the visual areas that support these behaviors or the development of non-visual regions downstream that receive the read-out from areas showing early arealization.

Our approach for measuring the organization of visual cortex in human infants is to perform retinotopic mapping with functional magnetic resonance imaging (fMRI) — the gold standard for defining visual areas V1–V4 in older children and adults (Conner, Sharma, Lemieux, and Mendola, 2004; Wandell, Dumoulin, and Brewer, 2007). Indeed, no prior study has used experimental stimuli to define the boundaries and properties of V1–V4 in infant primates — work at this age in non-human primates relied on functional connectivity rather than on more standard mapping stimuli (Arcaro and Livingstone, 2017) and the youngest evidence of retinotopy in humans so far has come from 5 year olds (Conner, Sharma, Lemieux, and Mendola, 2004; Gomez, Natu, Jeska, Barnett, and Grill-Spector, 2018). The reason why this was not previously performed in human infants is that fMRI studies in awake infants (human or non-human) present many challenges. Some of these challenges are general, such as head motion and fussiness, whereas others are specifically problematic for retinotopic mapping, such as an inability to instruct or enforce eye fixation (Braddick and Atkinson, 2011; Ellis and Turk-Browne, 2018). Moreover, the organization of infant visual cortex may not be detectable at the macroscopic level accessible to fMRI (Chapman, Gödecke, and Bonhoeffer, 1999). That said, there is some reason for optimism: fMRI has revealed that infant visual cortex is functionally interconnected (Gao, Lin, Grewen, and Gilmore, 2017), shows evoked responses to visual inputs (Ellis, Skalaban, Yates, Bejjanki, Córdova, and Turk-Browne, 2020a), and responds in a localized way to motion (Biagi, Crespi, Tosetti, and Morrone, 2015) and categories (Deen, Richardson, Dilks, Takahashi, Keil, Wald, Kanwisher, and Saxe, 2017). Nonetheless, it is unknown whether infants have retinotopic maps, which has long been considered critical for grounding the study of infant vision (Braddick and Atkinson, 2011).

To perform retinotopic mapping, we used a new protocol that enables fMRI in awake and behaving infants (Ellis, Skalaban, Yates, Bejjanki, Córdova, and Turk-Browne, 2020a,b; Ellis, Skalaban, Yates, and Turk-Browne, 2020c). In individual infants from 5 to 23 months old, we sought to define ventral and dorsal V1, V2, and V3, dorsal V3A/B, and ventral V4. In an alternating block design, we used meridian mapping (horizontal vs. vertical) to identify area boundaries between quarter-field (V1–V3) and half-field (V3A/B, V4) representations (Fox, Miezin, Allman, Van Essen, and Raichle, 1987; Schneider, Noll, and Cohen, 1993) and spatial frequency mapping (high vs. low) as a proxy for foveal and peripheral representations of eccentricity within these areas (Arcaro and Livingstone, 2017; Henriksson, Nurminen, Hyvärinen, and Vanni, 2008). We chose these stimuli because they are relatively more tolerant to inconsistent fixation than traveling wave (Engel, Rumelhart, Wandell, Lee, Glover, Chichilnisky, and Shadlen, 1994; Sereno, Dale, Reppas, Kwong, Belliveau, Brady, Rosen, and Tootell, 1995) or population receptive field (Dumoulin and Wandell, 2008) approaches (Wandell, Dumoulin, and Brewer, 2007). Namely, the desired stimulation was received wherever the infant fixated on the stimulus. These two measures of arealization provided independent, potentially divergent, measures of functional maturity in infant visual cortex. This allowed us to test whether arealization is present early in infancy, as well as how it develops over our age range.

## Results

We showed 17 infants (5–23 months old) blocks of stimuli (Figure 1) for meridian mapping (Fox, Miezin, Allman, Van Essen, and Raichle, 1987; Schneider, Noll, and Cohen, 1993) and spatial frequency mapping (Arcaro and Livingstone, 2017; Henriksson, Nurminen, Hyvärinen, and Vanni, 2008). In meridian mapping blocks, large horizontally and vertically oriented meridian bow ties were shown for 20 s each in a counterbalanced order. In spatial frequency mapping blocks, low and high spatial frequency Gaussian random fields were shown for 20 s each in a counterbalanced order. Throughout all blocks, an illustrated movie was shown at center to encourage fixation. Blocks were separated by 6 s of rest.

**Figure 1:**
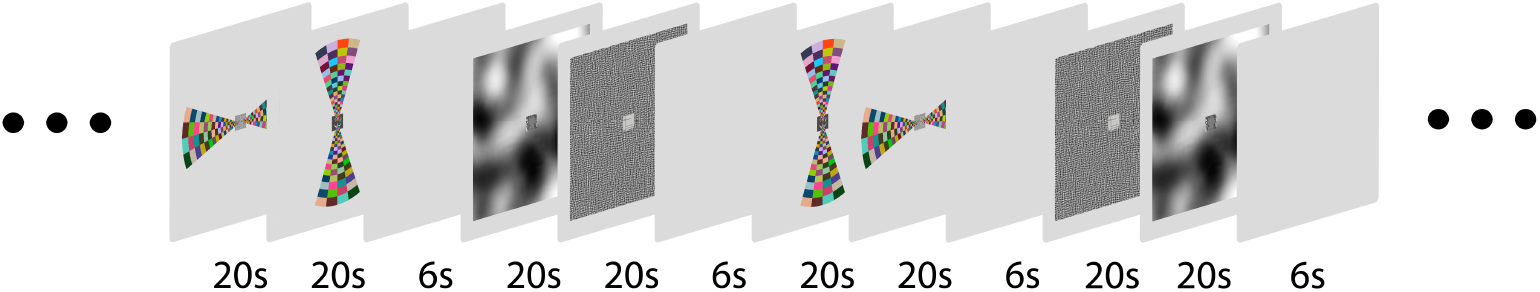
Experimental stimuli for retinotopic mapping. Participants viewed wide-field (40° visual angle) meridian or spatial frequency stimuli on the ceiling of the scanner bore in an alternating block design. Each block contained two phases in a counterbalanced order: vertical and horizontal for meridian mapping, high and low for spatial frequency mapping. The two phases lasted 20 s each and appeared back-to-back with no break, followed by 6 s of rest. A small illustrated movie (1.5°) was played at center to encourage fixation.

### Looking behavior

Gaze was manually coded from a video recording of the infant’s face during the scan. In meridian mapping blocks, participants similarly looked at the horizontal meridian (*M*=0.92) and vertical meridian (*M*=0.92) for almost the entirety of the block (difference CI=[−0.02, 0.02], *p*=0.866). As expected, and motivating our use of task designs tolerant to eye movements, looking at the meridian did not necessarily mean fixating the movie at center. Infants often looked left or right of center on horizontal meridians (proportion *M*=0.41) and above or below center on vertical meridians (*M*=0.30). There was a trend toward more off-center looking for horizontal vs. vertical (difference CI=[0.000, 0.221], *p*=0.050), which may reflect the ease of saccading in azimuth compared to elevation (Aslin and Salapatek, 1975). In spatial frequency blocks, participants similarly looked at high spatial frequencies (*M*=0.90) and low spatial frequencies (*M*=0.91) for almost the entirety of the block (difference CI=[−0.040, 0.014], *p*=0.310). Infants infrequently looked off-center on high spatial frequencies (*M*=0.08) and low spatial frequencies (*M*=0.05). There was reliably more off-center looking for high vs. low (difference CI=[0.003, 0.070], *p*=0.033), which may reflect a need to foveate high-frequency stimuli (given that the movie was shown at fixation). Overall, stimulation was uniform in the visual field the majority of the time, and the remaining time was still usable because of designs that ensured similar stimulation regardless of where the stimulus was fixated.

### Volumetric analyses

A general linear model (GLM) was used to estimate blood oxygenation level-dependent (BOLD) responses to the four stimulus conditions. As a sanity check, infant visual cortex was robustly activated when collapsing across conditions (Figure 2a). Indeed, contrasts between horizontal and vertical meridians (Figure 2b) and between low and high spatial frequencies (Figure 2c) in each participant’s volumetric space indicate strong differential responding to the conditions.

**Figure 2:**
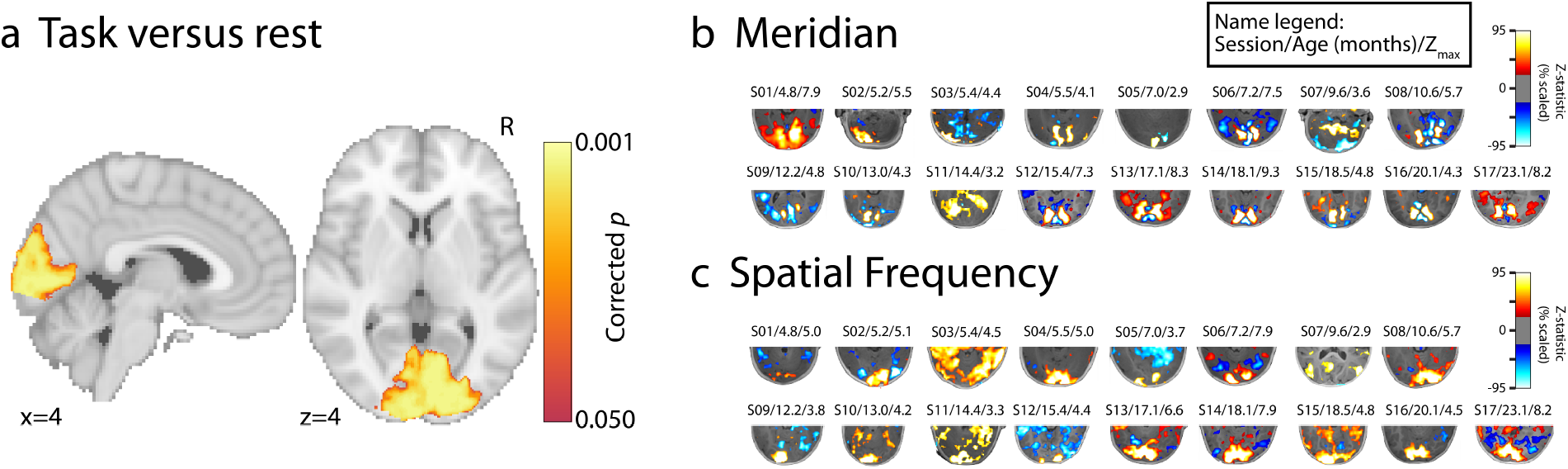
Statistical maps in volumetric space. (a) Significant voxels (TFCE corrected *p*<0.05) from an *F*-test across participants, demonstrating reliable evoked BOLD activity in standard space. (b) Contrast of horizontal greater than vertical meridians on each participant’s anatomical image for the posterior portion of the brain. The minimum *Z*-statistic threshold is 1.96 and the maximum differs by participant, indicated by Z_max_. Participants are ordered youngest to oldest from top left to bottom right. The text above each volume indicates: session number/age in months/Z_max_. (c) Analogous contrast of high greater than low spatial frequency. Refer to Table S1 for details about the participants.

### Evidence of arealization

To determine whether infants have retinotopic organization, we first created surface reconstructions using iBEAT v2.0 (Li, Nie, Wang, Shi, Gilmore, Lin, and Shen, 2014; Li, Wang, Shi, Gilmore, Lin, and Shen, 2015; Li, Wang, Yap, Wang, Wu, Meng, Dong, Kim, Shi, Rekik, et al., 2019; Wang, Li, Shi, Cao, Lian, Nie, Liu, Zhang, Li, Wu, et al., 2018) (Figure S1). These surfaces were inflated and cut to make flatmaps. The contrast between horizontal and vertical meridians was projected onto each participant’s flatmap and used for tracing visual areas. Areas were traced based on the alternations in sensitivity to horizontal and vertical meridians using a suitable protocol for adults (Wandell, Dumoulin, and Brewer, 2007). For instance, the V1/V2 border was based on the peak in the vertical meridian representation on the gyral banks of the calcarine and the V2/V3 border was based on the next peak in the horizontal meridian representation. Figure 3a shows this contrast for an example 5.5-month old participant. Overlaid on this surface are the manually traced areas demarcating low- and mid-level visual cortex. In this participant we traced V1, V2, and V3 in both hemispheres and in both ventral and dorsal cortex, as well as left dorsal V3A/B and bilateral ventral V4. Although there was variability across participants (Figure 3b), differential sensitivity to horizontal and vertical meridians was clear enough to identify areas in 16 out of 17 infants (average of 6.6 out of 8 areas in each hemisphere).

**Figure 3:**
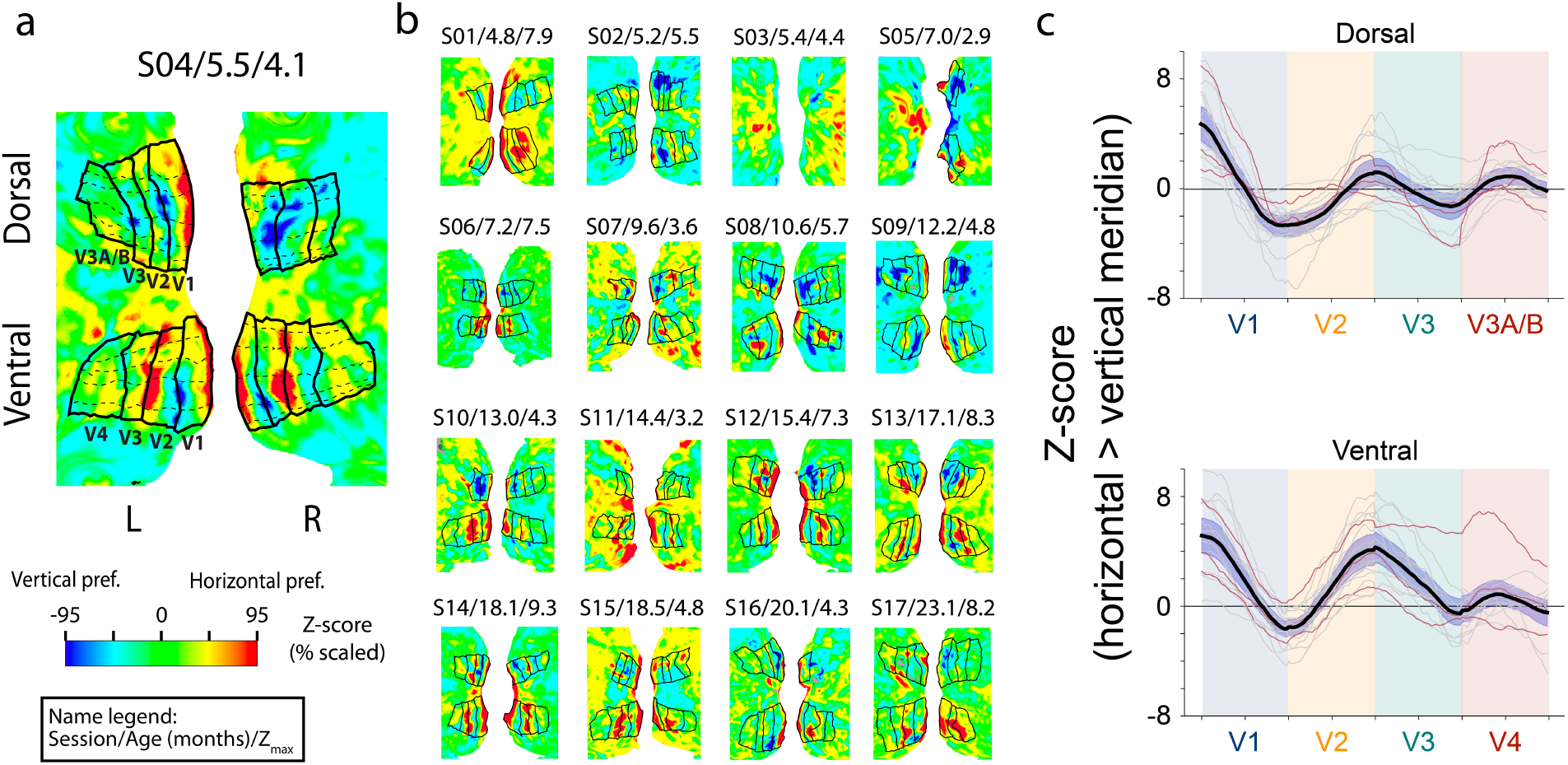
Arealization of infant visual cortex. (a) Example 5.5 month old participant with areas traced on the cortical surface. Labels shown for left V1, V2, V3, V3A/B and V4. Colors indicate the *Z*-statistic value for the contrast of horizontal greater than vertical meridian, scaled to the range of the 95th percentile of voxel *Z*-statistics in the occipital lobe (here, 4.1). Dashed lines drawn perpendicular to area boundaries were used to measure oscillations in sensitivity to horizontal and vertical meridians across areas. The text above each surface indicates: session number/age in months/Z_max_. (b) Statistical maps for all 16 other participants, ordered youngest to oldest from top left to bottom right. Refer to Table S1 for details about the participants. (c) Contrast values of horizontal greater than vertical meridian for points on lines drawn perpendicular to the area boundaries, separately for dorsal and ventral areas. The extracted values were interpolated to a normalized length across areas. The black line indicates the average of all participants, the gray lines indicate participants over 6 months, and the red lines indicate participants under 6 months. The purple shaded region around the black line is the 95% confidence interval across participants.

To verify that there were gradients of selectivity to different meridians across the visual areas (Arcaro, McMains, Singer, and Kastner, 2009), we traced lines perpendicular to the area boundaries, starting at the fundus of the calacarine sulcus and progressing anterior, and extracted the contrast values from points along those lines (dashed lines on Figure 3a). There were reliable oscillations in sensitivity to horizontal and vertical meridians across areas (Figure 3c). Although this analysis is not suitable for statistical analysis (the same data were used to trace the areas), it remains a useful analysis to quantify the amount of change across areas. The difference between the peaks for vertical and horizontal meridians was smaller in anterior compared to posterior areas, potentially reflecting the larger receptive fields and thus coarser retinotopic maps of these anterior areas. Furthermore, this analysis shows the consistency of these patterns across participants. Indeed, three participants under 6 months also showed this same pattern, providing evidence of early retinotopic organization.

To evaluate the reliability of our manual tracings, we examined whether the resulting areas were more similar within versus between infants in the subset of four participants with two or three sessions. Participant surfaces were aligned to standard space and the similarity of the manually traced areas was computed between participants using the Dice coefficient. Similarity was higher within the same participant across sessions (*M*=0.47) than between different participants yoked to have the same age difference (*M*=0.35, CI=[0.08, 0.17], *p*<0.001). In fact, when a participant was compared across sessions, they were often more similar to themselves than to almost any other participant (rank *M*=2.0, *p*<0.001). This was true for even the youngest participant with two sessions (S02 at 5.2 months and S05 at 7.0 months). Such reliability of the functional topographies is remarkable because the surfaces were based on anatomical scans from different ages that could vary in size and cortical folding.

### Spatial frequency tuning

We next examined sensitivity to eccentricity in infant visual cortex. Standard eccentricity mapping requires presenting stimuli at different distances from fixation. Because we could not instruct or enforce fixation, such stimuli are not viable in infants. Instead, we used spatial frequency tuning as a proxy for eccentricity (Arcaro and Livingstone, 2017; Henriksson, Nurminen, Hyvärinen, and Vanni, 2008; Tolhurst and Thompson, 1981): neurons sensitive to high spatial frequencies have small receptive fields that tend to process close eccentricities near the fovea and that are more prevalent in low-level visual areas; conversely, neurons sensitive to low spatial frequencies have larger receptive fields that tend to process farther eccentricities in the periphery and that are more prevalent in mid- and high-level visual areas. Thus, to assess voxel-wise sensitivity to eccentricity, we used a GLM to contrast BOLD activity for high vs. low spatial frequency conditions. Figure 4a shows the spatial frequency map from a 5.5-month old participant. There is a gradient from the fovea within each area, as well as the gradient from low- to mid-level visual cortex. This pattern was observed across all participants with traced areas (Figure 4b).

**Figure 4:**
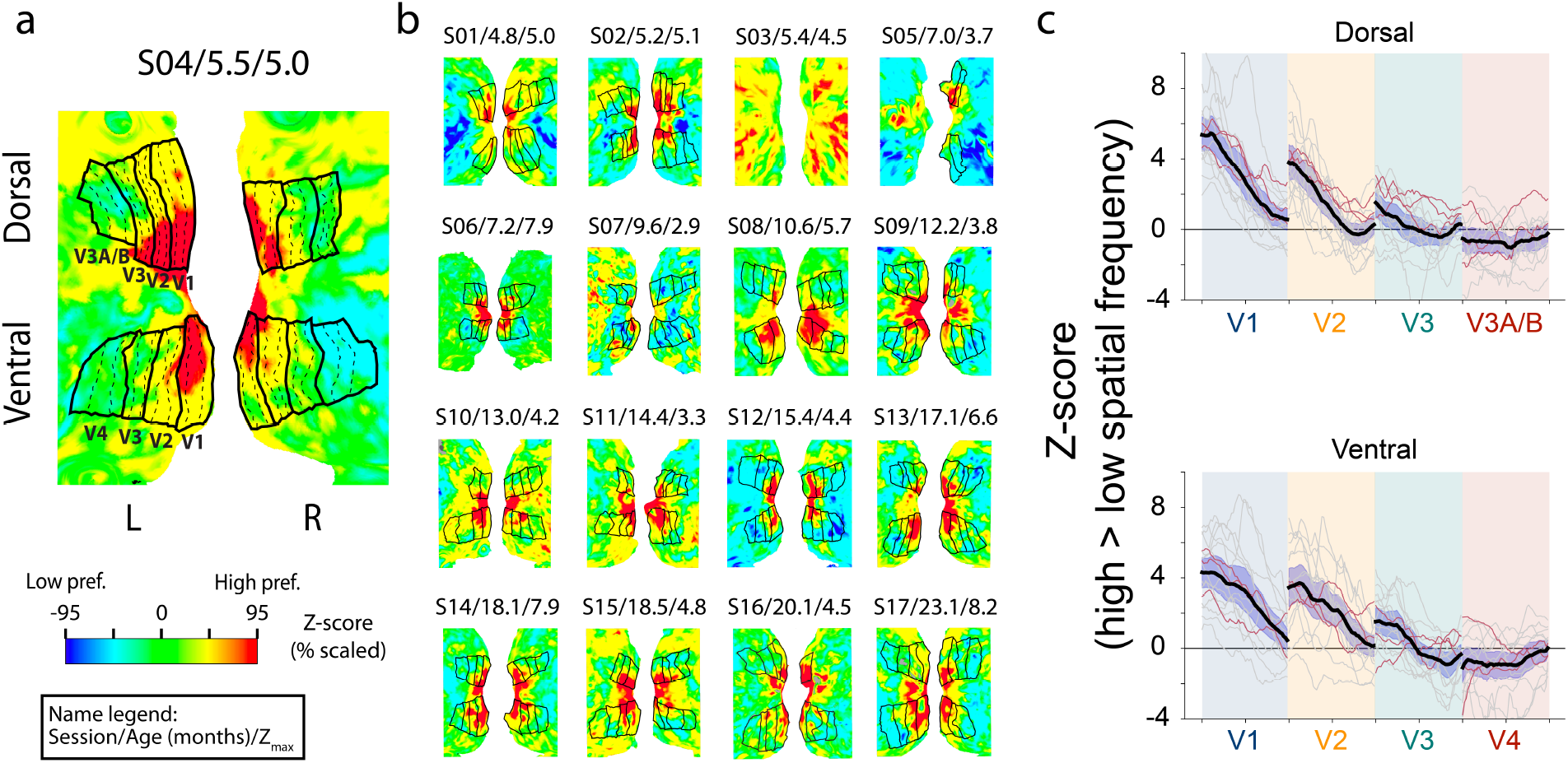
Spatial frequency tuning in infant visual cortex. (a) Example 5.5 month old participant with areas traced on the cortical surface. Colors indicate the *Z*-statistic value for the contrast of high greater than low spatial frequency, scaled to the range of the 95th percentile of voxel *Z*-statistics in the occipital lobe (here, 5.0). Dashed lines drawn parallel to area boundaries were used to measure gradients in sensitivity to high and low spatial frequencies across areas. The text above each surface indicates: session number/age in months/Z_max_. (b) Statistical maps for all 16 other participants, ordered youngest to oldest from top left to bottom right. Refer to Table S1 for details about the participants. (c) Contrast values of high greater than low spatial frequency for points on lines drawn parallel to the area boundaries, separately for dorsal and ventral areas. Each area is demarcated by a colored column. The foveal boundary of the area is on the left side of the column. The extracted values were interpolated to a normalized length across areas. The black line indicates the average of all participants, the gray lines indicate participants over 6 months, and the red lines indicate participants under 6 months. The purple shaded region around the black line is the 95% confidence interval across participants.

To quantify the gradient in sensitivity to spatial frequency within areas, we traced lines with each area parallel to the area boundaries (dashed lines on Figure 4a). Importantly, these areas were drawn using the meridian mapping blocks, so the tracings were independent from the spatial frequency data. We measured the contrast of high greater than low spatial frequency along the lines, starting at the foveal boundary. Sensitivity to spatial frequency transitioned significantly from foveal (first quarter of the line) to peripheral (last quarter of the line) edges (Figure 4c), in dorsal and ventral V1 (CI=[3.18, 4.55], *p*<0.001), V2 (CI=[2.53, 4.22], *p*<0.001), and V3 (CI=[0.96, 2.25], *p*<0.001), but not V4 (CI=[−1.23, 0.05], *p*=0.081) or V3A/B (CI=[−0.96, 0.61], *p*=0.695). This confirms a gradient in sensitivity to spatial frequency from the fovea to periphery in infant visual cortex, including in infants under 6 months. Indeed, the first vs. last quarter difference did not reliably correlate with age in V1 (*r*=0.00, *p*=0.958), V2 (*r*=0.04, *p*=0.793), V4 (*r*=0.26, *p*=0.258), or V3A/B (*r*=0.18, *p*=0.520), although was stronger in older children in V3 (*r*=0.52, *p*=0.014).

We also observed a gradient in sensitivity to spatial frequency across areas, with a greater overall response to high vs. low spatial frequency in V1 vs. V2 (CI=[0.82, 1.50], *p*=<0.001), V2 vs. V3 (CI=[1.21, 2.10], *p*<0.001), ventral V3 vs. V4 (CI=[0.38, 1.51], *p*=0.001), and dorsal V3 vs. V3A/B (CI=[0.52, 1.26], *p*<0.001). This confirms differences in spatial frequency tuning between low- and mid-level areas in infant visual cortex, as has been observed in the adult visual cortex (Henriksson, Nurminen, Hyvärinen, and Vanni, 2008). This same pattern was present in our participants under 6 months. In fact, the average difference between sensitivity to high vs. low spatial frequency did not reliably correlate with age in V1 (*r*=0.35, *p*=0.162), V2 (*r*=0.27, *p*=0.300), V3 (*r*=−0.12, *p*=0.571) V3A/B (*r*=−0.39, *p*=0.140), or V4 (*r*=−0.13, *p*=0.683). These findings are consistent with the possibility that gradients of spatial frequency tuning within and across areas are largely stable across infancy.

### Area configuration and size

The results so far fit an early arealization hypothesis, in which area boundaries and spatial frequency tuning are established in infants as young as 5 months. However, this does not preclude the possibility of hierarchical maturation in other properties of these areas, including their configuration and size. To test for such age effects, we extended the Dice similarity analysis. Rather than compare maps within individual infants across repeat sessions, here we compare the manual tracings of areas V1–V4 in standard space across different infants (Figure 5a). We first tested whether maps are more similar for participants of similar ages, which would lead to a negative correlation between age difference and Dice coefficient (closer in age = higher similarity). However, this relationship was not reliable and in fact was numerically in the wrong direction (*r*=0.11, *p*=0.247). We then tested another potential age effect in which older infants are more similar to each other than younger infants are to each other. This would lead to a positive correlation between the average age of the two infants whose maps are being compared and their Dice coefficient (older age = higher similarity). We excluded repeat sessions from this analysis because they occurred more often in older infants and yielded higher Dice coefficients, and thus could bias the correlation. Across the remaining pairs of sessions between participants (N = 114), we observed a robust positive correlation (Figure 5b; *r*=0.36, *p*<0.001). This correlation could be explained if we were better able to align older infants into standard space. However, the number of manually coded alignment errors (Thompson, Stein, Medland, Hibar, Vasquez, Renteria, Toro, Jahanshad, Schumann, Franke, et al., 2014) was not significantly correlated with age across sessions (*r*=−0.39, *p*=0.185). The alignment of visual gyri and sulci between each participant and standard space also was not significantly related to age across sessions (*r*=0.14, *p*=0.609).

**Figure 5:**
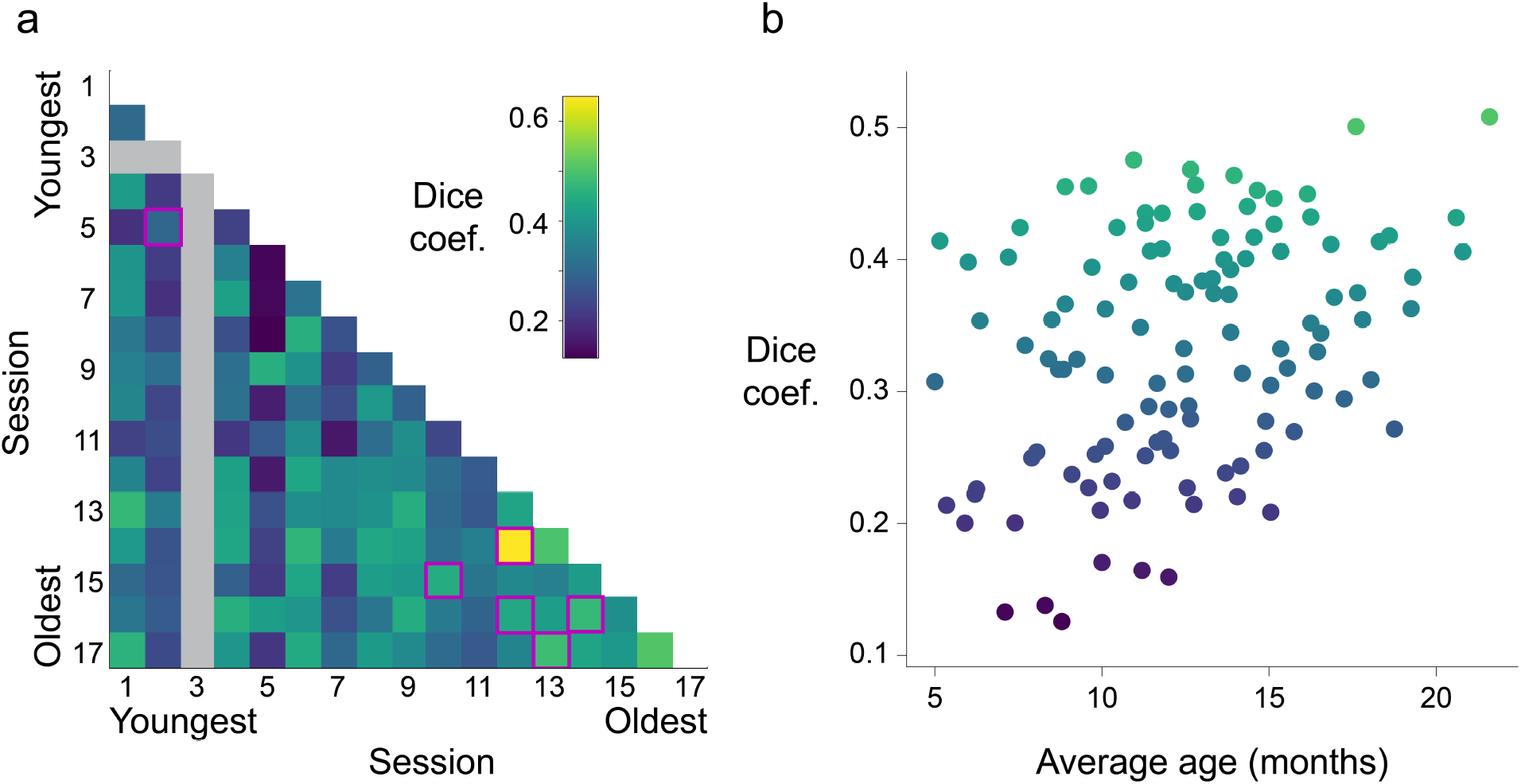
Overlap of retinotopic maps across infants. After alignment to standard space, the cortical locations of the manually traced visual areas were compared across pairs of sessions using the Dice coefficient. (a) Rows and columns are ordered from youngest to oldest, with each cell depicting a session pair. The purple boxes highlight repeat sessions from the same participants. The gray strip corresponds to a participant (S03) with no traced visual areas. (b) Correlation of Dice similarity with the average age (in months) of the infants in the two sessions being compared. Note that repeat sessions from the same infant are not visualized because they were excluded from the analysis to avoid biasing in favor of a correlation.

We interpret the increased similarity of older infants as reflecting convergence toward mature retinotopic organization. To evaluate this, we computed the Dice coefficient between the manual tracings of infant areas V1–V4 and a comprehensive atlas of adult visual areas in standard space (Wang, Mruczek, Arcaro, and Kastner, 2014). Overall, the similarity of infants to the standard adult atlas was high (*M*=0.39). In fact, infants were more similar to this atlas than to other infants (*M*=0.33, CI=[0.03, 0.08], *p*<0.001). There was an overall age effect, with similarity to the adult atlas increasing as a function of infant age (Figure 6; *r*=0.41, *p*=0.042). This is qualified by the fact that similarity did not increase systematically within infants across repeat sessions. Indeed, the high similarity even for the youngest infants suggests that any maturation rests on a foundation of early, adult-like arealization. The high similarity is likely also an underestimate given differences in the visual extent of the stimuli between adults (30° visual angle diameter) and infants (40°), and greater coverage of the fovea in the adult atlas. Moreover, different approaches to retinotopy were used for adults (traveling wave) and infants (meridian). The fact that the adult atlas still provided such a good guide to infant visual areas further helps validate the alignment of infant data to standard space, and suggests that such atlases could plausibly be used as a starting point for ROIs in future studies.

**Figure 6:**
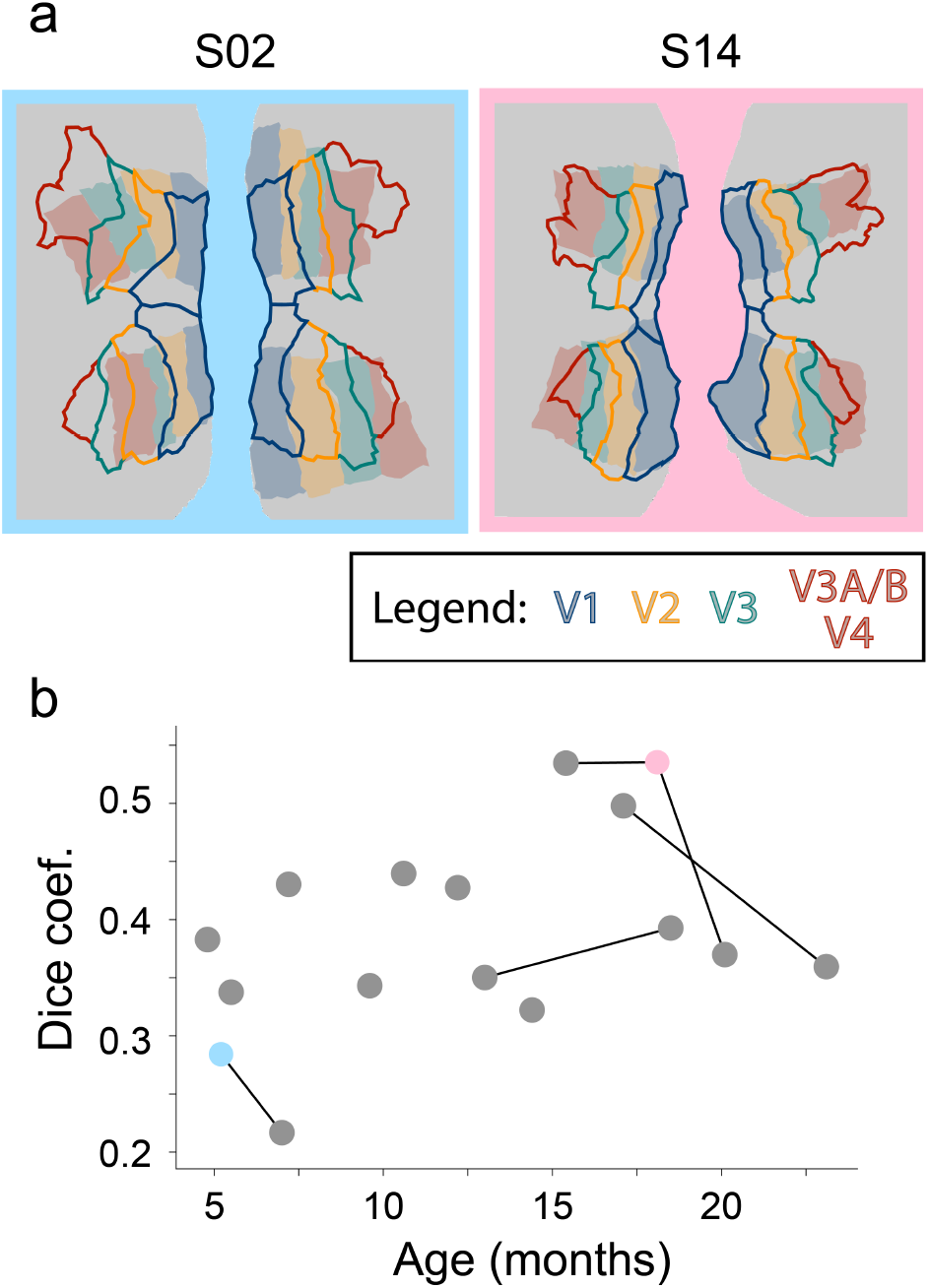
Overlap of infant visual areas with an adult atlas. Participants were aligned into standard space and compared to the relevant areas from an adult atlas (Wang, Mruczek, Arcaro, and Kastner, 2014). (a) Two example infants with their manually traced visual areas in shaded fill colors and the corresponding visual areas from the standard space atlas overlaid in color outlines. (b) Dice coefficient for each participant between visual areas from manual tracing in infants and standard atlas in adults. Lines connect the same participant across sessions. Cyan and pink dots correspond to participants S02 and S14, respectively.

An alternative way to evaluate region maturation is to consider how the size of each area changes with age. The hierarchical maturation hypothesis would predict that low-level areas are mature in size by early infancy, at least relative to other areas, whereas the size of mid- and high-level areas would change over development. The most direct measure of size is surface area; however, this metric is imprecise near the foveal boundary because of the video we played at fixation. We therefore used the length between area boundaries (dashed lines in Figure 3a) as our measure of size (although obtained similar results for surface area; Figure S2). Figure 7 shows the relationships across the 16 participants with traceable areas. There was a significant relationship between size and age in V1 (*r*=0.78, *p*<0.001) and V2 (*r*=0.50, *p*=0.013), the relationship was marginal in V3 (*r*=0.54, *p*=0.059), and not significant in V3A/B (*r*=0.18, *p*=0.517) or V4 (*r*=0.23, *p*=0.398). These changes in V1 and V2 size could reflect the global growth of the brain over age in our sample (*r*=0.80, *p*<0.001). However, the relationship between size and age persisted after controlling for global volume in V1 (partial *r*=0.57, *p*=0.024), though not V2 (*r*=0.29, *p*=0.208) (Table S2). Hence, we see that only the earliest visual areas change in size across early development, counter to what would be expected from the perspective of hierarchical maturation of visual cortical areas.

**Figure 7:**
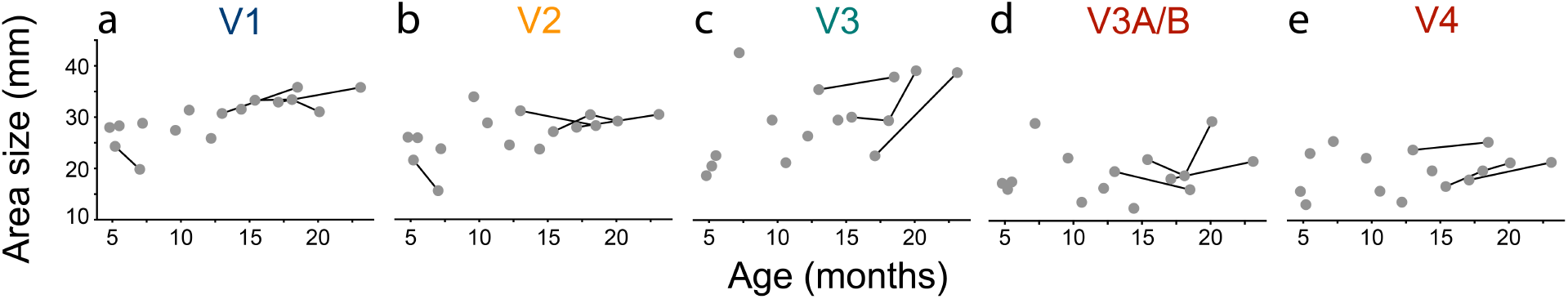
Relationship between the size of visual areas and infant age. Size is operationalized as the length between area boundaries averaged over hemispheres, summed over ventral and dorsal streams (for V1, V2, and V3). For a comparable analysis of surface area, see Figure S2. Lines are used to connect data from the same participant tested more than once across sessions. Raw data for each area are reported in Table S2.

## Discussion

We investigated the presence of retinotopic organization in human infants as young as 5 months. We found evidence of visual areas V1–V4 even in our youngest participants and these areas were reliable within participant across sessions. Moreover, there was a gradient in sensitivity from high to low spatial frequency within visual areas as well as a gradient across areas in the hierarchy, matching the topography of eccentricity and spatial frequency tuning in adults (Henriksson, Nurminen, Hyvärinen, and Vanni, 2008). Together, these results support the existence of a hierarchical, retinotopic organization across visual cortex early in development. Area boundaries and spatial frequency tuning did not consistently vary with age in our sample, although the size of V1 (but not the other areas) did increase with age controlling for global brain growth. The weak evidence for change in retinotopic organization over this age range suggests that the development of visual cortex from 5–23 months may reflect fine-tuning rather than reorganization.

We examined two properties of the visual system to evaluate the hierarchical maturation hypothesis that visual areas develop sequentially from low-, to mid-, to high-level areas (Bourne and Rosa, 2006; Condé, Lund, and Lewis, 1996; Distler, Bachevalier, Kennedy, Mishkin, and Ungerleider, 1996; Gomez, Natu, Jeska, Barnett, and Grill-Spector, 2018; Kiorpes, 2016; Zhang, Zheng, Watanabe, Maruko, Bi, Smith, and Chino, 2005). First, we examined the presence and development of visual areas by defining their boundaries with meridian mapping. Our evidence that visual areas exist even early in infancy is consistent with anatomical findings from non-human primates (Bourne and Rosa, 2006). However, functional confirmation of visual areas has not been previously reported in human infants. Even if these areas are established, the hierarchical maturation hypothesis might suggest that properties like area size would develop sequentially. Although we did observe age-related changes in the size of visual areas, they were strongest in low-level areas (V1 and V2), contrary to what this hypothesis would predict. Second, we tested for gradients in spatial frequency tuning within and across areas with spatial frequency mapping. Gradients in preference from high to low spatial frequencies are thought to reflect changes in receptive field size from smaller to larger, respectively (Arcaro and Livingstone, 2017; Henriksson, Nurminen, Hyvärinen, and Vanni, 2008). Mirroring changes that occur with eccentricity, there were spatial frequency gradients from the fovea to the periphery within visual areas and overall from low- to mid-level visual areas. There was reason to believe that gradients might develop at these levels of the hierarchy in infancy, given that the receptive field properties of high-level areas change later in childhood (Gomez, Natu, Jeska, Barnett, and Grill-Spector, 2018). However, we observed gradients in low- and mid-level areas even in our youngest participants, and the strength of these gradients was not reliably related to age.

Considering the two lines of evidence together, our results are consistent with an early arealization account where arealization is present and stable in infancy for low- and mid-level visual cortex. Critically, hierarchical maturation of the visual system could be occurring in human development but outside of the age range tested here, either earlier in neonates/fetuses or later in toddlers or children. Additionally, more research is needed to explore how other components of visual cortical function develop in infancy in these areas. Even so, our results suggest that important architectural features of the visual system are established early in development. One consequence is that gradients in receptive field size may be available to scaffold the development of functionally selective high-level visual cortex (Arcaro, Schade, Vincent, Ponce, and Livingstone, 2017; Gomez, Barnett, and Grill-Spector, 2019; Hasson, Levy, Behrmann, Hendler, and Malach, 2002). In particular, this architecture may enable high-level visual cortex to develop stereotyped localization of category-selective cortex (e.g., faces require high precision near fovea, buildings require precision in the periphery; (Hasson, Levy, Behrmann, Hendler, and Malach, 2002)). Our results also suggest that substantial changes in visual processing during infancy (Baker, Norcia, and Candy, 2011; Kovacs, Kozma, Feher, and Benedek, 1999; Lewis and Maurer, 2005; Patel, Maurer, and Lewis, 2010; Siu and Murphy, 2018) do not hinge on the emergence of retinotopic organization in low- or mid-level visual cortex. One possibility is that improvements in visual processing over infancy and childhood could relate to improvements in downstream regions responsible for the read-out of initial visual cortical processing, like high-level visual, association, or frontal cortices (Kiorpes and Movshon, 2014).

The protocol we developed for drawing areas was modeled on adult retinotopy (Arcaro, McMains, Singer, and Kastner, 2009), which has been extensively validated with animal models (Wandell, Dumoulin, and Brewer, 2007). A potential limitation of our study is that these comparisons may not be appropriate for human infants. For instance, tracing human V4 is notoriously difficult, in part because of the venous eclipse (Winawer, Horiguchi, Sayres, Amano, and Wandell, 2010) and in part because it is hard to distinguish the anterior border of V4 from surrounding areas like VO1. These challenges may be exacerbated in infants, for example, because their areas are smaller and their scans are poorer quality. On the other hand, area boundaries in infant visual cortex, even if imprecise, were reliable within and between participants in our sample.

We used a block-design approach that is uncommon in adult retinotopy. Meridian mapping designs like ours have drawbacks (Wandell, Dumoulin, and Brewer, 2007), such as limited coverage of the visual field and potentially mislabeling voxels that are only partially responsive to a stimulus. However, our goal was to demarcate visual areas (Kastner, De Weerd, and Ungerleider, 2000; Schneider, Noll, and Cohen, 1993; Shipp, Watson, Frackowiak, and Zeri, 1995; Tootell, Reppas, Kwong, Malach, Born, Brady, Rosen, and Belliveau, 1995), rather than estimate the selectivity of individual voxels (cf. population receptive fields; (Dumoulin and Wandell, 2008)), so these drawbacks are less pertinent. Indeed, the block design we used is relatively robust because it can survive the exclusion of a small number of individual time-points if the participant looks away or moves their head excessively. This is a key advantage for infant fMRI, in which data loss is expected within blocks. More critically, the traditional traveling wave approach (Engel, Rumelhart, Wandell, Lee, Glover, Chichilnisky, and Shadlen, 1994) requires central fixation throughout each block. In our experience, this is impractical in human infants. Fixation cannot be instructed or enforced, and they look away from center approximately one third of the time, even with a movie at fixation. That said, if a traveling wave approach could somehow be adapted to these constraints, this might enable tracing of high-level visual areas (Arcaro, McMains, Singer, and Kastner, 2009) that cannot be resolved with meridian mapping (Kastner, De Weerd, and Ungerleider, 2000; Shipp, Watson, Frackowiak, and Zeri, 1995; Tootell, Reppas, Kwong, Malach, Born, Brady, Rosen, and Belliveau, 1995).

In sum, we found robust evidence for retinotopic organization in awake behaving human infants, ranging in age from 5 to 23 months old. We identified four areas in both ventral and dorsal streams, spanning low- and mid-level visual cortex, and these areas were reliable across sessions. The receptive field properties within and across the areas mirrored what has been observed in adults. We found limited evidence of developmental change in retinotopic organization, other than the size of low-level areas. Together, these results suggest that the early infant visual system has the basic cortical architecture needed for low- and mid-level visual processing. These results also provide a foundation for understanding infant visual abilities, such as acuity, contrast sensitivity, and contour integration, as well as the neural basis of infant visual dysfunction in disorders such as amblyopia (Braddick and Atkinson, 2011). More broadly, retinotopy gave credibility to fMRI research during its earliest days by validating that architectural features of visual cortex first identified in animals could be measured in human BOLD (Engel, Rumelhart, Wandell, Lee, Glover, Chichilnisky, and Shadlen, 1994; Schneider, Noll, and Cohen, 1993; Sereno, Dale, Reppas, Kwong, Belliveau, Brady, Rosen, and Tootell, 1995). Likewise, the current study shows the promise of fMRI with awake behaving infants to reveal the function of the infant brain and to track changes across early postnatal development.

## Acknowledgements

Thank you to N. Wilson and N. Córdova for help with initial experiment design. Thank you to K. Armstrong, L. Rait, J. Daniels and the entire Yale Baby School team for recruitment, scheduling, and administration. Thank you to A. Bracher for help with scanning. Thank you to J. Fel, and J. Wu for help with gaze coding. Thank you to K. Anderson for help with the iBEAT to Freesurfer conversion. Thank you to R. Watts for technical support. We are grateful for internal funding from the Department of Psychology and Princeton Neuroscience Institute at Princeton University and from the Department of Psychology and Faculty of Arts and Sciences at Yale University. N.B.T-B. was further supported by the Canadian Institute for Advanced Research.

## Author Contributions

C.T.E., M.J.A., & N.B.T-B. initially created the experiment design. C.T.E., L.J.S., V.R.B., & N.B.T-B. did early piloting. C.T.E., T.S.Y., & N.B.T-B. collected the data. C.T.E. & T.S.Y. preprocessed the data. C.T.E. & M.J.A performed the analyses. All authors contributed to the drafting of the manuscript.

## Methods

### Participants

Data from 17 sessions with infants aged 4.8 to 23.1 months (*M*=12.2, SD=5.7; 13 females) met our inclusion criteria of at least one usable phase per condition. The planned sample size was to have two participants in each three-month window between 3 and 24 months. We met this criteria, except we only had one participant between 21 and 24 months. This sample size is larger than previous developmental studies of retinotopy (Conner, Sharma, Lemieux, and Mendola, 2004). Not included in the sample are data from 15 sessions with enough blocks prior to exclusions for head motion and eye gaze, or from four sessions without enough blocks even prior to exclusions. The final sample included 12 unique participants, three participants who provided two sessions of usable data, and one participant who provided three sessions. These sessions occurred at least one month apart (range=1.8–6.0) and so the data from these sessions were treated separately, similar to prior work (Deen, Richardson, Dilks, Takahashi, Keil, Wald, Kanwisher, and Saxe, 2017). Refer to Table S1 for information on each participant. Data was collected at the Brain Imaging Center (BIC) at Yale University. Parents provided informed consent on behalf of their child. The study was approved by the Human Subjects Committee at Yale University.

### Data acquisition

Data were acquired with a Siemens Prisma (3T) MRI with the bottom of the 20-channel Siemens head coil. Anatomical images were acquired with a T1-weighted PETRA sequence (TR_1_=3.32ms, TR_2_=2250ms, TE=0.07ms, flip a ngle=6°, m atrix=320×320, slices=320, resolution=0.94mm iso, radial slices=30000). For three of our younger, compliant participants, we also collected a T2-weighted SPACE sequence (TR=3200ms, TE=563ms, flip angle=120°, matrix=192×192, slices=176, resolution=1mm isotropic), and these were supplied to iBEAT to support surface reconstruction. Functional images were acquired with a whole-brain T2* gradient-echo EPI sequence (TR=2s, TE=30ms, flip angle=71°, matrix=64×64, slices=34, resolution=3mm iso, interleaved slice acquisition).

### Data and code availability

The code for running the retinotopy task can be found at: https://github.com/ntblab/experiment_menu/tree/retinotopy/. The code for performing the analyses can be found at: https://github.com/ntblab/infant_neuropipe/tree/Retinotopy/. The data, including anonymized anatomical images, surface reconstructions, manually traced areas, and both raw and preprocessed functional images can be found at: (to be shared on Dryad when published).

### Procedures

There are many challenges when conducting fMRI research with awake infants. We have described and validated our approach in a separate methods paper (Ellis, Skalaban, Yates, Bejjanki, Córdova, and Turk-Browne, 2020a). In brief, families visited the lab before their first scanning session for an orientation session. The aim of this was to acclimate the infant and parent to the scanning environment. Scanning sessions were scheduled for a time when the parents thought that the infant would be calm. The infant and parent were extensively screened for any metal on or in their body. Hearing protection for the infant consisted of three layers: silicon inner ear putty, over-ear adhesive covers, and ear muffs. The infant was positioned on the scanner bed, on top of a vacuum pillow that reduced movement. The top half of the head coil was not used because the bottom elements provided sufficient coverage of the smaller infant head. This allowed for sufficient visibility to monitor the infant’s comfort and allowed us to project stimuli onto the ceiling of the bore directly above the infant’s face using a custom mirror system. A video camera (MRC high-resolution camera) allowed us to record the infant’s face during scanning for monitoring and offline eye gaze coding.

When the infant was focused, stimuli were shown in MATLAB using Psychtoolbox (http://psychtoolbox.org), against a gray background. For the meridian mapping blocks, a bow tie cut-out of a pastel-colored checkerboard was presented in either a vertical or horizontal orientation (Tootell, Reppas, Kwong, Malach, Born, Brady, Rosen, and Belliveau, 1995). The arcs of the bow ties were 45° and their diameter spanned 40 visual degrees. The checkerboard spacing increased logarithmically out from the fovea, approximating the cortical magnification factor (Tootell, Reppas, Kwong, Malach, Born, Brady, Rosen, and Belliveau, 1995). The color of the checkerboard alternated every 125ms between the original color pattern and its negative. For the spatial frequency mapping blocks, the stimuli were grayscale Gaussian random fields of high (1.5 cycles per visual degree) or low (0.05 cycles per visual degree) spatial frequency (Arcaro and Livingstone, 2017). The difference between these spatial frequencies should be detectable to 3-month old infants (Banks, Stephens, and Hartmann, 1985), although spatial frequency discrimination develops into adolescence (Patel, Maurer, and Lewis, 2010; van den Boomen and Peters, 2017). These patterns were shown as squares spanning 40° on each edge. There were five images for each spatial frequency, one of which was shown every 500 ms. Meridian and spatial frequency blocks contained two phases of stimulation. The first phase consisted of one of the conditions (e.g., horizontal or high) for 20s, followed immediately by the second phase with the other condition of the same block type (e.g., vertical or low, respectively) for 20s. The order of conditions was counter-balanced across blocks. At the end of each block there was at least 6s rest before the start of the next block. Participants alternated between blocks of the spatial frequency and meridian mapping tasks, with the goal of acquiring 8 blocks total.

To facilitate attention to the center of the stimulus, a movie was played in a small window (1.5° in diameter) at that location. For spatial frequency blocks this was overlaid on top of the stimulus, for meridian mapping blocks the bow tie was overlaid on the movie. The movie showed grayscale shapes moving in unpredictable patterns, including jittering, looming, and smooth motion. The movie was saturated in order to minimize the amount of high contrast changes.

### Gaze coding

Infant gaze was coded offline by two or three coders (*M*=2.12) blind to condition. The coders determined whether the eyes were oriented left, right, up, down, center, off-screen (i.e., blinking or looking away), or were undetected (i.e., out of the camera’s field of view). Codes for the directional gazes (i.e., left, right, up, and down) were only applied if the coder believed the infants were still looking at the screen. For instance, if the participant was looking left but off the screen then that frame was coded as off-screen. For the spatial frequency blocks, the gaze was considered acceptable if it was coded as center or any of the four directions. For the meridian mapping blocks, gaze directions perpendicular to the bow tie orientation were treated as equivalent to off-screen in preprocessing. For example, if the infant was viewing a horizontal bow tie, looking to the left, center, or right was considered acceptable.

The frame rate and resolution varied across participants, but the minimum rate was 16Hz and we always had sufficient resolution to identify the eye. The label for each frame was determined as the mode of a moving window of five frames centered on that frame across all coder reports. In case of a tie, we used modal response from the previous frame. The coders were highly reliable: when coding the same frame, coders reported the same response on 79% (range across participants=71– 86%) of frames. Phases were excluded if the participant was coded as looking away from the stimulus for more than 25% of the time, computed separately for the two phases within each block. This was a stricter criterion than our other infant fMRI studies (Ellis, Skalaban, Yates, Bejjanki, Córdova, and Turk-Browne, 2020a,b; Ellis, Skalaban, Yates, and Turk-Browne, 2020c) because of the importance of eye position for retinotopy. Across all included phases, participants looked at the stimulus 91% of the time on average (range across participants=87–95%).

To determine whether there were differences in the looking behavior across the conditions, we computed each participant’s average proportion of looking time for every manual code that was allowable in that phase (e.g., left, center, or right for horizontal meridian). We quantified the difference in mean looking for horizontal and vertical conditions using bootstrap resampling. We also compared the mean looking time away from center per condition (e.g., left or right for horizontal meridian) with bootstrap resampling.

### Preprocessing

Individual runs were preprocessed using FEAT in FSL (https://fsl.fmrib.ox.ac.uk/fsl), with optimizations for infant fMRI data. We discarded three volumes from the beginning of each run, along with the volumes automatically discarded by the EPI sequence. Blocks were stripped of any excess burn-in or burn-out volumes beyond the 3 TRs (6s) that were planned. Some runs contained other experiments not discussed here (N=15 sessions). In such cases, pseudo runs were created containing only the data of interest. Blocks were sometimes separated by long pauses (>30s) within a session because of a break, an anatomical scan, or an intervening experiment (N=5; *M*=556.5s break; range=107.9–1183.4s). We used the ‘centroid’ volume (i.e., with the minimal Euclidean distance from all other volumes) for alignment and motion correction. Slice-time correction was applied to realign the slices in each volume. Time-points were excluded if the head motion between time-points exceeded 3mm (average in blocks with at least one usable phase: *M*=1.9%, range=0.0–7.0%), and phases were excluded if more than 50% of TRs exceeded this threshold. We interpolated rather than removed these time-points in order to avoid biasing the linear detrending (in later analyses these time-points were removed). To make the mask of brain voxels, we thresholded the signal-to-fluctuating-noise ratio (SFNR) of each voxel in the centroid volume at the trough in the histogram of values. The data were smoothed with a Gaussian kernel (5mm FWHM) and linearly detrended. AFNI’s despiking algorithm attenuated aberrant time-points within voxels. To account for differences across runs in intensity and variance, the blocks that were considered usable were normalized over time using *Z*-scoring, prior to the runs being concatenated for further analyses. For further explanation and justification of this preprocessing procedure, please refer to (Ellis, Skalaban, Yates, Bejjanki, Córdova, and Turk-Browne, 2020a).

Participants were excluded if they did not have at least 1 usable phase from each of the four conditions. After these criteria were applied, participants (coincidentally) had a similar number of phases of each condition on average: 3.53 (SD=1.33, range: 1–6) high spatial frequency, 3.47 (SD=1.72, range: 1–6) low spatial frequency, 3.47 (SD=1.46, range: 1–6) horizontal meridian, and 3.47 (SD=1.46, range: 1–6) vertical meridian.

Each run’s centroid volume was registered to the infant’s anatomical scan from the same session. FLIRT with a normalized mutual information cost function was used for initial alignment. Additional manual registration was performed using mrAlign from mrTools (Gardner lab) to fix deficiencies of automatic registration. The preprocessed functional data were aligned into anatomical space with their original spatial resolution (3mm iso). This aligned data were mapped on to surface space, as described below. Whole-brain voxelwise analyses required further alignment of functional data into a standard space. For alignment to standard, the anatomical scan from each participant was automatically (FLIRT) and manually (Freeview) aligned to an age-specific MNI infant template (Fonov, Evans, Botteron, Almli, McKinstry, and Collins, 2011) and then aligned to the adult MNI template (MNI152). The functional data were transformed into standard space for the task vs. rest contrast. To determine which voxels to consider at the group level, the intersection of brain voxels from all infant participants in standard space was used as a whole-brain mask.

For surface reconstruction, we used iBEAT v2.0 to acquire the surfaces (Li, Nie, Wang, Shi, Gilmore, Lin, and Shen, 2014; Li, Wang, Shi, Gilmore, Lin, and Shen, 2015; Li, Wang, Yap, Wang, Wu, Meng, Dong, Kim, Shi, Rekik, et al., 2019; Wang, Li, Shi, Cao, Lian, Nie, Liu, Zhang, Li, Wu, et al., 2018). The output of the iBEAT pipeline is the inner and outer surfaces, as well as the volumetric segmentation of gray-matter and white-matter. Figure S1 shows the surface reconstructions overlaid on a slice of the anatomical data for each participant. These were then inserted into a FreeSurfer-style pipeline (a walkthrough is provided in the codebase). As part of the pipeline, these surfaces were inflated into spheres and aligned to the Buckner40 template (Dale, Fischl, and Sereno, 1999). To investigate the quality of the surfaces and the alignment to standard space, we used the ENIGMA consortium quality control procedure (Thompson, Stein, Medland, Hibar, Vasquez, Renteria, Toro, Jahanshad, Schumann, Franke, et al., 2014), in which we evaluated defects in the projection of the Desikan-Killany atlas onto the individual data. In particular, we quantified how many errors there were in each hemisphere for the following regions (atlas labels) that are known to be prone to poor segmentation: bankssts, precentral, postcentral, pericalcarine, parahippocampal, entorhinal, rostralanteriorcingulate, insula. The infant data showed typical amounts of errors compared to what is reported for adult data. The data were then resampled in SUMA, using an icosahedral shape, into standard space with a constant number of nodes (Argall, Saad, and Beauchamp, 2006). To generate flatmaps, a convexity map (based on the position of gyri and sulci) was computed using AFNI (Cox, 1996) and inflated brains were cut and flattened using the FreeSurfer procedure. Once flattened, statistics maps were projected on to the surface for evaluation.

### GLM analysis

For the main analyses, a GLM was fit to the BOLD activity in each voxel using FEAT in FSL. Two separate GLMs were performed, one containing the horizontal and vertical meridian regressors, and the other containing the high and low spatial frequency regressors. Each regressor modeled phases with a boxcar lasting 20s, convolved with a double-gamma hemodynamic response function. The six translation and rotation parameters from motion correction were included in the GLM as nuisance regressors. TRs that were excluded (i.e., had translational motion greater than 3mm) were scrubbed by including an additional regressor for each to-be-excluded time-point (Siegel, Power, Dubis, Vogel, Church, Schlaggar, and Petersen, 2014). The condition regressors were then contrasted to find the differential evoked response. The voxelwise *Z*-statistic volumes for these contrasts were extracted for each participant. For visualization purposes, we set the maximum value of these maps to be the 95th percentile of the *Z*-statistic value for each participant in a large, anatomically defined occipital mask.

To test for visual-evoked activity, an *F*-test was performed in FSL using each phase type (i.e., horizontal and vertical meridians, high and low spatial frequencies) as regressors in a GLM to identify which voxels respond to any visual stimulation. As a conservative test to evaluate where the brain was most activated across participants, we *Z*-scored the resulting *F*-values within each participant. Hence, any mean differences in *F*-values across participants are mitigated. Instead, what matters is whether the *F*-values are high, relative to other voxels in that participant, in the same voxels across participants. The resulting statistic map was volumetrically aligned to standard space and then merged across all participants. We used threshold free cluster enhancement through the randomise function in FSL, resulting in voxel clusters *p*<.05 corrected.

### Visual area and gradient tracing

The contrast maps for horizontal greater than vertical were transformed onto the flat surface map and used for tracing (Argall, Saad, and Beauchamp, 2006). Traditional tracing guidelines for adult humans were followed (Wandell, Dumoulin, and Brewer, 2007). The areas traced were ventral V1, V2, V3 (also known as VP), and V4 (also known as hV4), and dorsal V1, V2, V3 and V3A/B. The border between V1 and V2 was defined by the peak in the vertical meridian, the border between V2 and V3 was defined by the peak in the horizontal meridian, the border between V3 and ventral V4 or dorsal V3A/B was defined by the peak in the vertical meridian, and the terminal border of ventral V4 and dorsal V3A/B was defined by the peak of the vertical meridian after a half cycle. It is typical to trace only these areas using a meridian mapping paradigm (Kastner, De Weerd, and Ungerleider, 2000; Shipp, Watson, Frackowiak, and Zeri, 1995; Tootell, Reppas, Kwong, Malach, Born, Brady, Rosen, and Belliveau, 1995), as well as early traveling wave studies (Sereno, Dale, Reppas, Kwong, Belliveau, Brady, Rosen, and Tootell, 1995). This procedure is likely to lead to imprecision between the V4 boundary and the VO1 boundary, and did not provide sufficient resolution to demarcate V 3A and V 3B. When the distinctions between areas were not clear, the maximum range of the colormap was varied. If this did not resolve the ambiguity then the evoked response to just the vertical meridian (rather than the contrast between horizontal and vertical) was checked. If these additional steps did not clarify the boundary between areas, the area was not drawn. The peripheral extent of the area was estimated by the horizontal greater than vertical contrast and the *F*-test for each participant. Because of the movie shown at fixation, the foveal response was expected to be contaminated, hence the areas were not traced to the foveal confluence. To test how the foveal confluence varied with age, we also traced the areas that bridged between the ventral and dorsal areas of V1, V2, and V3 for each hemisphere.

To quantify alternations in sensitivity to horizontal vs. vertical meridians across visual areas (Figure 3c), we traced additional lines of interest in the visual cortex using SUMA (Figure 3a; (Arcaro, McMains, Singer, and Kastner, 2009)). These lines were drawn perpendicular to the area boundaries, posterior to anterior, for areas that were traced. Five lines, spaced equally along the width of the areas, were drawn for each hemisphere and the dorsal and ventral areas. These lines were drawn using only the areal boundaries for guidance, making the coder unaware of the local intensity changes within areas. Nodes in surface space on those lines were indexed for their values of the horizontal greater than vertical meridian contrast. The number of nodes along each line varied between areas and participants. In order to standardize the size of these lines for the sake of comparison, we interpolated the values along each line to contain 50 values within each area (up to 200 total).

The gradients in sensitivity from high to low spatial frequency (Figure 4c) were quantified by tracing lines within each area, parallel to the areal boundaries (Figure 4a). Two lines were traced from the foveal to the peripheral boundary of each area and were used to index the values of the high greater than low spatial frequency contrast. The indexed values along these lines were also interpolated to include 50 values.

To statistically test the gradients in sensitivity from high to low spatial frequency within area, we divided the lines running parallel to the area boundary into quartiles, with the first quartile containing the section of the line closest to the foveal confluence. The contrasts for ventral and dorsal areas were averaged for lines in V1, V2, and V3. We averaged the contrast of high greater than low spatial frequency for the first quartile and compared it to the fourth quartile using bootstrap resampling (Efron and Tibshirani, 1986). Namely, we sampled, with replacement, the difference in contrast values from all participants 10,000 times, averaging across participants on each iteration to generate a sampling distribution. Confidence intervals reflect the 2.5 and 97.5 percentiles of this distribution. For null hypothesis testing, we calculated the *p*-value as the proportion of samples whose mean was in the opposite direction from the true effect, doubled to make the test two-tailed. To test whether the difference in contrast between quartiles varied with age, we used bootstrap resampling of the correlation, randomly sampling bivariate data from 17 participants with replacement and calculating the Pearson correlation on each of 10,000 iterations. We calculated the *p*-value as the proportion of samples resulting in a correlation with the opposite sign from the true correlation, doubled to make the test two-tailed.

To statistically test the gradient of spatial frequency tuning across (rather than within) areas, we averaged the contrast values along the whole lines and compared adjacent areas (i.e., V1 vs. V2, V2 vs. V3, etc.) using bootstrap resampling. Areas were averaged across dorsal and ventral, except when comparing ventral V3 with V4 and dorsal V3 with V3A/B. We used bootstrap resampling of the correlation to test the relationship between age and the averaged contrast within area.

### Comparing visual areas

The consistency of the traced visual areas across participants was quantified using the Dice coefficient (Dice, 1945). This was computed by finding every node in the surface that was labeled as belonging to a visual area and comparing whether those nodes had the same label across participants. The Dice coefficient is a fraction where the number of matching nodes, multiplied by 2, is the numerator, and the total number of nodes with a label in either participant is the denominator. For a given pair of sessions, the Dice coefficient was quantified for each hemisphere separately and averaged. Only areas that were traced in both participants were considered. This means the denominator of the Dice coefficient is not inflated by labels that could not possibly match between participants. The Dice coefficient was calculated for all pairwise comparisons between sessions. Comparisons of the same participant across multiple sessions are used to determine whether there is greater reliability over time within vs. between participants. To approximately match the ages of the participants being compared, we started with one of the participants with repeat sessions and calculated the age differential; we then found another participant with the most similar age difference to one of these sessions (mean difference in age between the repeat session and the matched participant = 0.97 months). For example, consider the participant who was 5.2 months at their first session (S02) and 7.0 months at their next session (S05). To provide a between-participant control, we found the participant (S06) who was closest in age to the second session (7.2 months). We then compared the Dice coefficient of S02 and S05 with the Dice coefficient of S02 and S06. To evaluate this statistically, we used bootstrap resampling of the difference in Dice coefficients between the within- and between-participant pairs. The Dice coefficient comparing a participant to themselves across sessions was ranked against the Dice coefficients from relating that participant to all other participants. To evaluate significance of this rank, for each participant with a repeat session we generated a random rank in the possible range of values for that participant and averaged across the group. We did this 10,000 times to get a distribution of permuted ranks. We then compared the observed average rank to this permuted distribution to find the likelihood of finding a ranking as extreme as the one we observed.

To test how similarity between participants varied with age, we compared the ages of the participants to their Dice coefficient. We first tested whether similarity in age predicted the Dice coefficient. To do this, we subtracted the ages of participants and related the absolute value of this difference to the Dice coefficient. We next tested if the overall age of the participants predicted the Dice coefficient. To do this, we compared the average age of each pair of participants with the Dice coefficient for each pair. Bootstrap resampling was used to evaluate the significance of these correlations. In both of these analyses, comparisons using the same participant across multiple sessions were ignored because the sampling of these participants was skewed to older ages. Because within-participant Dice coefficients were higher, this skew would have biased the correlations to be positive.

To measure the quality of alignment to the standard template, we used two complementary metrics. First, we quantified the number of defects in the projection of the Desikan-Killany atlas, as described above. Second, we correlated the convexity map of individual participants with the convexity map from standard space (from fsaverage (Dale, Fischl, and Sereno, 1999)). The convexity maps of the individual and standard surface were masked according to the relevant regions (V1–V4) that were labelled in the atlas of visual cortex (Wang, Mruczek, Arcaro, and Kastner, 2014). The correlation between the convexity map and standard space was computed for each hemisphere separately. This correlation was Fisher transformed and then averaged between hemispheres. Bootstrap resampling of the correlation was used to quantify the significance of the relationship between age and the measure of alignment.

To determine the degree to which the visual areas have adult-like localization, the Dice coefficient was calculated between the traced areas in each infant and those same areas as defined in an atlas of the visual hierarchy (Wang, Mruczek, Arcaro, and Kastner, 2014). All of the areas we traced were available for comparison in the atlas except for V3A/B, which is separated in the atlas but was combined here. The maximum probability surface was used so that each node was uniquely assigned to a specific area, if at all.

The Dice coefficient to the standard atlas was computed for each individual and related to their age to determine whether the similarity to adults changes across early development. The significance of this correlation was computed using bootstrap resampling. We also tested whether the Dice coefficient was higher between two participants than between one participant and the atlas. The Dice coefficients comparing each participant with the other participants (excluding comparisons across sessions from the same participant) were averaged and subtracted from the Dice coefficient from that participant and the atlas. This difference for each participant was then tested for significance with bootstrap resampling.

Our main measure of the visual area size was the length of each traced area. This length was computed on surfaces in native space by taking all pairwise distances of nodes along the lines traced running perpendicular to the area boundaries and averaging them (Figure 3a). That is, length corresponds to the distance between boundaries. Length was chosen as the primary metric over other measurements like surface area because it is less sensitive to challenges in drawing the foveal border. Nevertheless, a similar pattern of results was obtained with surface area, computed by averaging the extent of the white matter and pial surface for all nodes labeled as belonging to an area. For V1, V2, and V3, we first averaged the lengths across hemispheres and then added these averages for the ventral and dorsal areas. This was still possible if an area was traced in only the left or right hemisphere, but if either the ventral or dorsal areas had not been traced in both the left and right hemisphere then the length of this area was not estimated. The average length of the remaining participants was related to their age using bootstrap resampling. To observe size changes independent of overall brain size, a partial correlation was computed where overall brain size (as measured based on the skullstripped volume from iBEAT v2.0) was used as a covariate. Analyses using gray matter volume as a covariate produced similar results. This partial correlation was also evaluated statistically using bootstrap resampling. To test whether the foveal confluence varied in size with age, the area of each foveal area was divided by the area of the sum of dorsal and ventral area for that area, and then correlated with age using bootstrap resampling.

## Supplementary Information

**Figure S1:**
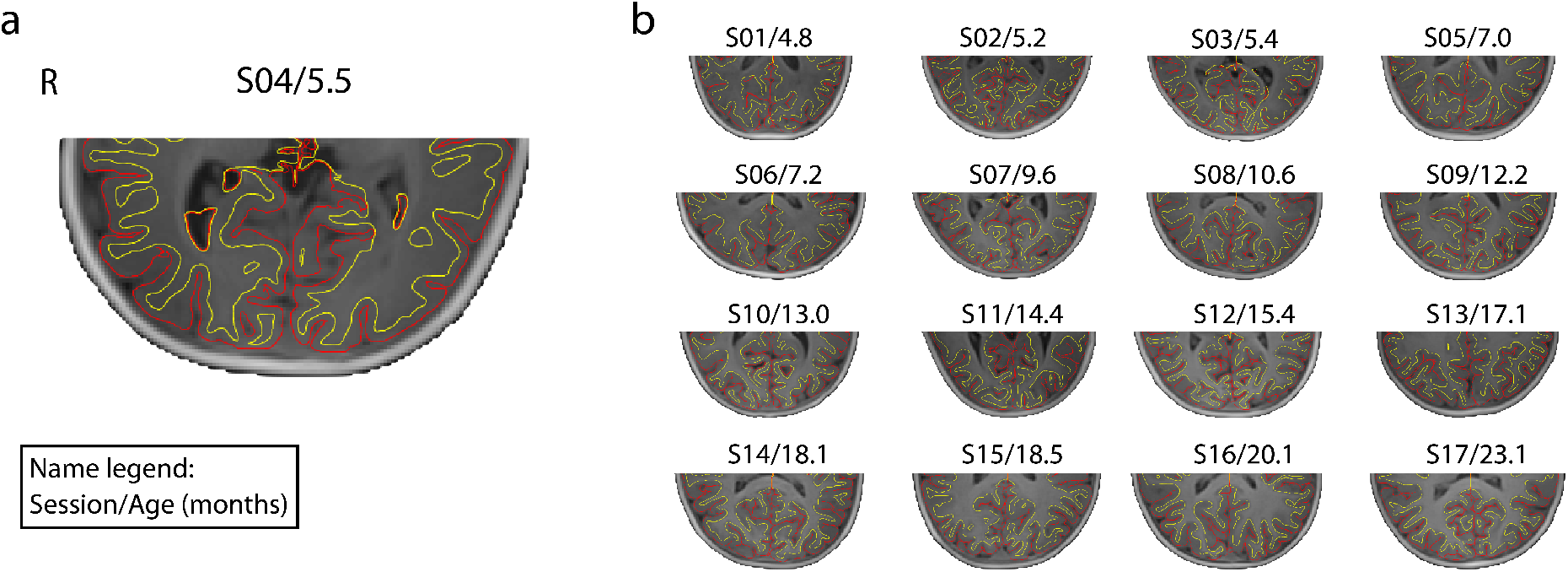
Pial and white matter surfaces from iBEAT v2.0. The posterior portion of anatomical images with surfaces outlined for (a) an example 5.5-month old participant, and (b) all other participants. The yellow line is the white matter surface and the red line is the pial surface. The text above each volume indicates: session number/age in months. No editing was performed on any of these surfaces before they were inflated.

**Figure S2:**
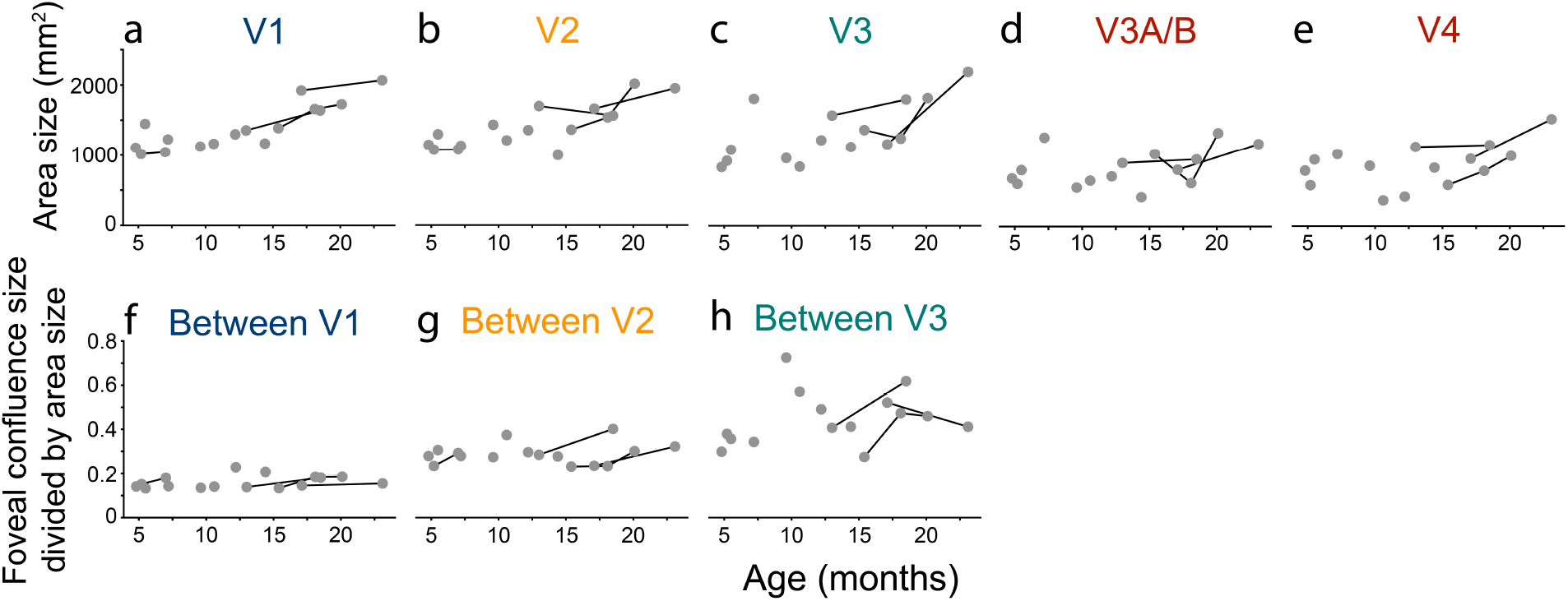
Relationship between surface area and the participant’s age, akin to Figure 7. The size was first averaged across hemispheres. For V1, V2, and V 3, ventral and dorsal areas were summed. Lines connect the same participant across sessions. Raw data for each area are reported in Table S3. There were strong positive correlations with age in V1 (*r*=0.84, *p*=0.001), V2 (*r*=0.79, *p*<0.001), and V3 (*r*=0.66, *p*=0.016), but not V3A/B (*r*=0.39, *p*=0.170) or V4 (*r*=0.46, *p*=0.088). Controlling for whole-brain volume, the partial correlation of area size and age remained reliable in V1 (*r*=0.79, *p*=0.014) and V2 (*r*=0.66, *p*=0.021), but not in V3 (*r*=0.36, *p*=0.476), V3A/B (*r*=0.24, *p*=0.432), or V4 (*r*=0.26, *p*=0.610). Moreover, imprecision in mapping area boundaries near the fovea (potentially caused by poor fixation) did not seem to confound the tests of agerelated changes in area size: the surface area of the unmapped visual cortex around the fovea, relative to the area of the mapped dorsal and ventral areas, showed a marginal effect in V1 (*r*=0.32 *p*=0.054), but did not change with age in V2 (*r*=0.15 *p*=0.559) or V3 (*r*=0.24 *p*=0.282).

**Table S1:** Participant information for included participants. ‘Session num.’ is the session number used in figures to designate participants. ‘ID’ is a unique infant identifier (i.e., sXXXX Y Z), with the first four digits (XXXX) indicating the family, the fifth digit (Y) the child number within family, and the sixth digit (Z) the session number with that child. This is used in the data release for labelling participants. ‘Age’ is recorded in months. ‘Sex’ is female or male. ‘Vert. phases’ is the number of vertical meridian phases included. ‘Hori. phases’ is the number of horizontal meridian phases included. ‘Low phases’ is the number of low spatial frequency phases included. ‘High phases’ is the number of high spatial frequency phases included. ‘Prop. incl. TRs’ is the proportion of usable TRs from the included blocks. ‘Prop. incl. eye’ is the proportion of usable eye data from the included phases. ‘IRR’ is the inter-rater reliability for the gaze coding.

| Session num. | ID | Age | Sex | Vert. phases | Hori. phases | Low phases | High phases | Prop. incl. TRs | Prop. incl. eye | IRR |
| --- | --- | --- | --- | --- | --- | --- | --- | --- | --- | --- |
| S01 | s2077_1_1 | 4.8 | M | 5 | 4 | 1 | 3 | 1.00 | 0.92 | 0.76 |
| S02 | s2097_1_1 | 5.2 | M | 4 | 4 | 4 | 4 | 1.00 | 0.95 | 0.83 |
| S03 | s8047_1_1 | 5.4 | F | 2 | 3 | 1 | 2 | 0.97 | 0.93 | 0.80 |
| S04 | s4047_1_1 | 5.5 | F | 3 | 3 | 4 | 4 | 1.00 | 0.95 | 0.82 |
| S05 | s2097_1_2 | 7.0 | M | 1 | 1 | 4 | 2 | 0.99 | 0.94 | 0.81 |
| S06 | s7017_1_3 | 7.2 | F | 6 | 6 | 6 | 6 | 1.00 | 0.95 | 0.78 |
| S07 | s7047_1_1 | 9.6 | F | 1 | 1 | 1 | 1 | 1.00 | 0.91 | 0.77 |
| S08 | s7067_1_4 | 10.6 | F | 3 | 3 | 2 | 3 | 1.00 | 0.89 | 0.78 |
| S09 | s8037_1_2 | 12.2 | F | 2 | 2 | 1 | 3 | 0.90 | 0.89 | 0.75 |
| S10 | s4607_1_4 | 13.0 | F | 5 | 5 | 4 | 6 | 0.93 | 0.87 | 0.72 |
| S11 | s1607_1_4 | 14.4 | M | 3 | 2 | 3 | 4 | 0.94 | 0.89 | 0.79 |
| S12 | s6687_1_4 | 15.4 | F | 5 | 6 | 6 | 4 | 1.00 | 0.91 | 0.84 |
| S13 | s8687_1_5 | 17.1 | F | 4 | 4 | 4 | 2 | 0.98 | 0.89 | 0.86 |
| S14 | s6687_1_5 | 18.1 | F | 5 | 4 | 6 | 5 | 1.00 | 0.89 | 0.84 |
| S15 | s4607_1_7 | 18.5 | F | 2 | 3 | 3 | 4 | 1.00 | 0.92 | 0.71 |
| S16 | s6687_1_6 | 20.1 | F | 4 | 3 | 4 | 4 | 0.97 | 0.92 | 0.83 |
| S17 | s8687_1_8 | 23.1 | F | 4 | 5 | 5 | 3 | 0.98 | 0.91 | 0.76 |
| Av. | . | 12.19 | . | 3.47 | 3.47 | 3.47 | 3.53 | 0.98 | 0.91 | 0.79 |

**Table S2:** Length of visual areas per participant. ‘Session num.’ is the session number used in figures to designate participants. ‘ID’ is a unique infant identifier. ‘v’ before an area name denotes ventral. ‘d’ before an area name denotes dorsal. Length is measured in mm and is averaged across the left and right hemispheres, when available. The ‘V1’, ‘V2’ and ‘V3’ columns sum across the ventral and dorsal areas. ‘Volume’ is the size of the whole-brain mask in mm^3^.

| Session num. | ID | vV1 | vV2 | vV3 | V4 | dV1 | dV2 | dV3 | V3A/B | V1 | V2 | V3 | Volume |
| --- | --- | --- | --- | --- | --- | --- | --- | --- | --- | --- | --- | --- | --- |
| S01 | s2077_1_1 | 9.1 | 11.9 | 7.8 | 15.7 | 18.9 | 14.1 | 10.8 | 17.3 | 28.0 | 26.0 | 18.7 | 619858 |
| S02 | s2097_1_1 | 12.1 | 11.6 | 10.2 | 13.1 | 12.2 | 9.9 | 10.4 | 16.1 | 24.3 | 21.6 | 20.5 | 784387 |
| S03 | s8047_1_1 | nan | nan | nan | nan | nan | nan | nan | nan | nan | nan | nan | 739231 |
| S04 | s4047_1_1 | 13.9 | 14.0 | 12.6 | 23.1 | 14.4 | 11.9 | 9.9 | 17.6 | 28.3 | 25.9 | 22.6 | 797135 |
| S05 | s2097_1_2 | 8.4 | 10.5 | nan | nan | 11.5 | 5.1 | nan | nan | 19.8 | 15.6 | nan | 760968 |
| S06 | s7017_1_3 | 14.3 | 12.8 | 25.2 | 25.4 | 14.6 | 11.0 | 17.5 | 29.0 | 28.8 | 23.8 | 42.6 | 828623 |
| S07 | s7047_1_1 | 10.7 | 12.6 | 15.7 | 22.1 | 16.8 | 21.3 | 13.8 | 22.2 | 27.4 | 33.9 | 29.5 | 867040 |
| S08 | s7067_1_4 | 17.1 | 15.2 | 12.6 | 15.7 | 14.3 | 13.6 | 8.6 | 13.6 | 31.4 | 28.8 | 21.2 | 771459 |
| S09 | s8037_1_2 | 14.6 | 13.7 | 12.9 | 13.6 | 11.3 | 10.8 | 13.4 | 16.4 | 25.9 | 24.5 | 26.4 | 834345 |
| S10 | s4607_1_4 | 15.6 | 15.7 | 22.4 | 23.7 | 15.2 | 15.5 | 13.1 | 19.6 | 30.7 | 31.2 | 35.4 | 959789 |
| S11 | s1607_1_4 | 18.4 | 13.8 | 13.7 | 19.7 | 13.1 | 9.9 | 15.8 | 12.4 | 31.5 | 23.7 | 29.5 | 940908 |
| S12 | s6687_1_4 | 16.1 | 14.1 | 15.4 | 16.6 | 17.2 | 13.0 | 14.6 | 22.0 | 33.3 | 27.1 | 30.1 | 951723 |
| S13 | s8687_1_5 | 16.4 | 14.3 | 12.8 | 17.9 | 16.5 | 13.7 | 9.7 | 18.2 | 32.9 | 28.0 | 22.5 | 863904 |
| S14 | s6687_1_5 | 18.2 | 15.1 | 14.5 | 19.7 | 15.2 | 15.3 | 14.9 | 18.8 | 33.4 | 30.4 | 29.4 | 952569 |
| S15 | s4607_1_7 | 16.0 | 15.3 | 22.5 | 25.3 | 19.8 | 12.9 | 15.4 | 16.1 | 35.8 | 28.3 | 37.9 | 1045534 |
| S16 | s6687_1_6 | 14.5 | 18.4 | 16.7 | 21.2 | 16.6 | 10.3 | 22.4 | 29.4 | 31.0 | 28.8 | 39.1 | 978040 |
| S17 | s8687_1_8 | 20.2 | 15.2 | 21.4 | 21.3 | 15.6 | 15.3 | 17.4 | 21.6 | 35.8 | 30.5 | 38.8 | 913503 |
| Av. | . | 14.72 | 14.01 | 15.76 | 19.60 | 15.18 | 12.74 | 13.85 | 19.35 | 29.91 | 26.74 | 29.61 | 859354 |

**Table S3:** Surface area of visual areas per participant. ‘Session num.’ is the session number used in figures to designate participants. ‘ID’ is a unique infant identifier. ‘v’ before an area name denotes ventral. ‘d’ before an area name denotes dorsal. Area is measured in mm^2^ and is averaged across the left and right hemispheres, when available. The ‘V1’, ‘V2’ and ‘V3’ columns sum across the ventral and dorsal areas. ‘Volume’ is the size of the whole-brain mask in mm^3^.

| Session num. | ID | vV1 | vV2 | vV3 | V4 | dV1 | dV2 | dV3 | V3A/B | V1 | V2 | V3 | Volume |
| --- | --- | --- | --- | --- | --- | --- | --- | --- | --- | --- | --- | --- | --- |
| S01 | s2077_1_1 | 431 | 582 | 484 | 789 | 675 | 562 | 351 | 682 | 1106 | 1144 | 835 | 619858 |
| S02 | s2097_1_1 | 540 | 577 | 470 | 581 | 476 | 498 | 454 | 604 | 1016 | 1075 | 924 | 784387 |
| S03 | s8047_1_1 | nan | nan | nan | nan | nan | nan | nan | nan | nan | nan | nan | 739231 |
| S04 | s4047_1_1 | 681 | 711 | 609 | 948 | 765 | 584 | 474 | 800 | 1446 | 1294 | 1083 | 797135 |
| S05 | s2097_1_2 | 511 | 654 | nan | nan | 532 | 425 | nan | nan | 1043 | 1079 | nan | 760968 |
| S06 | s7017_1_3 | 592 | 557 | 937 | 1026 | 634 | 571 | 872 | 1264 | 1226 | 1128 | 1809 | 828623 |
| S07 | s7047_1_1 | 539 | 587 | 569 | 857 | 583 | 843 | 395 | 552 | 1122 | 1430 | 964 | 867040 |
| S08 | s7067_1_4 | 656 | 645 | 415 | 365 | 507 | 562 | 426 | 651 | 1163 | 1208 | 841 | 771459 |
| S09 | s8037_1_2 | 779 | 753 | 630 | 419 | 519 | 600 | 589 | 712 | 1299 | 1353 | 1219 | 834345 |
| S10 | s4607_1_4 | 705 | 864 | 1024 | 1127 | 650 | 833 | 547 | 903 | 1354 | 1697 | 1571 | 959789 |
| S11 | s1607_1_4 | 739 | 572 | 503 | 832 | 427 | 425 | 618 | 414 | 1167 | 997 | 1121 | 940908 |
| S12 | s6687_1_4 | 748 | 688 | 660 | 585 | 636 | 672 | 705 | 1031 | 1384 | 1360 | 1365 | 951723 |
| S13 | s8687_1_5 | 998 | 902 | 718 | 959 | 925 | 757 | 442 | 806 | 1922 | 1659 | 1161 | 863904 |
| S14 | s6687_1_5 | 1030 | 856 | 646 | 786 | 628 | 679 | 597 | 615 | 1658 | 1535 | 1243 | 952569 |
| S15 | s4607_1_7 | 841 | 820 | 1058 | 1154 | 796 | 740 | 741 | 955 | 1638 | 1561 | 1800 | 1045534 |
| S16 | s6687_1_6 | 1017 | 1272 | 856 | 1000 | 709 | 742 | 965 | 1329 | 1726 | 2014 | 1820 | 978040 |
| S17 | s8687_1_8 | 1043 | 930 | 1211 | 1524 | 1023 | 1019 | 980 | 1174 | 2067 | 1949 | 2191 | 913503 |
| Av. | . | 740.69 | 748.12 | 719.30 | 863.46 | 655.32 | 656.97 | 610.36 | 832.78 | 1396.02 | 1405.10 | 1329.67 | 859354 |

